# Peripheral Natural Killer cells from chronic hepatitis B patients display molecular hallmarks of T cell exhaustion

**DOI:** 10.1101/2020.06.16.154419

**Authors:** M. Marotel, M. Villard, I. Tout, L. Besson, O. Allatif, M. Pujol, Y. Rocca, M. Ainouze, G. Roblot, S. Viel, M. Gomez, V. Loustaud, S. Alain, D. Durantel, T. Walzer, U. Hasan, A. Marçais

## Abstract

A significant proportion of individuals infected by HBV develops chronic infection. Antiviral effectors such as Natural Killer (NK) cells have impaired functions in these patients, but the molecular mechanism responsible for this dysfunction remains poorly characterized. Here, we show that peripheral NK cells from chronic hepatitis B (CHB) patients have a defective capacity to produce IFN-γ, MIP1-β and TNF-α but retain an intact killing capacity. This functional phenotype was associated with a decrease in the expression of NKp30 and CD16, combined with defects in IL-15 stimulation of the mTOR pathway. Transcriptome analysis of NK cells in CHB patients further revealed a strong enrichment for transcripts typically expressed in exhausted T cells suggesting that NK cell dysfunction and T cell exhaustion rely on common molecular mechanisms. In particular, the transcription factor thymocyte selection-associated HMG box protein (TOX) and several of its targets, including immune checkpoints, were over-expressed in NK cells of CHB patients. This T cell exhaustion signature was predicted to be dependent on the calcium (Ca^2+^)-associated transcription factor NFAT. In line with this, when stimulating the Ca^2+^-dependent pathway in isolation, we recapitulated the dysfunctional phenotype. Thus, deregulated Ca^2+^ signalling could be a central event in both T cell exhaustion and NK cell dysfunction that occur during chronic infections.

## Introduction

Hepatitis B Virus (HBV) infection results in immune mediated viral clearance in 90-95% of adults. The remaining 5-10% fail to control viral infection due to failure in type 1 cellular immunity thus leading to chronic hepatitis B (CHB)^1^. As a result, more than 250 million individuals worldwide are chronic HBV carriers^2^. NK cells are endowed with antiviral properties such as IFN-γ and TNF-α secretion as well as cytotoxicity that could contribute to HBV clearance^3^. In patients, NK cells are activated during acute HBV infection before the onset of adaptive immunity^4–8^. However, in the chronic phase of the disease, NK cells present impaired functions, which could contribute to viral persistence. Indeed, NK cells from CHB patients are characterized by a decreased capacity to produce cytokines such as IFN-γ and TNF-α despite maintaining or even increasing their cytotoxic capacity^9–12^. This phenomenon has been termed “functional dichotomy”^9^. NK cell functions are controled by the relative strength of positive and negative signals triggered by the ligation of activating or inhibitory receptors^13^. In HBV infected patients, this balance might be skewed as a decrease in the expression of several activating receptors was previously observed in patient’s NK cells^7,11,12,14^. Furthermore, NK cells from CHB patients with high viral loads or liver damage display increased expression of certain inhibitory receptors including immune-checkpoint (ICP) markers such as NKG2A or T cell immunoglobulin and mucin domain containing 3 (TIM3)^15,16^. Moreover, cytokines also participate positively or negatively in the control of NK cell functions^17–20^. Molecularly, we and others have shown that the mechanistic target of rapamycin (mTOR) pathway integrates positive and negative signals derived from cytokines such as IL-15, IL-12 or TGF-β respectively to control NK cell metabolism and functions^18–21^. In the context of CHB, reports also involved immuno-modulatory cytokines such as IL-10 and TGF-β in the emergence/maintenance of the dysfunctional phenotype^10,11,22^. However, despite these pieces of evidence, a molecular framework to explain NK cell dysfunction is still missing. This is in contrast to the situation that prevails in the T cell field where dysfunction is also observed in contexts of persistent stimulation encountered during chronic infection or cancer: a phenomenon termed exhaustion. Indeed, T cell exhaustion has been defined as a stepwise differentiation process arising in situations of chronic stimulation and combining i) a gradual loss of effector functions and proliferative capacities, ii) an altered metabolism, iii) the expression of defined ICPs functioning as inhibitory receptors and iv) a specific transcriptional and epigenetic program^23^. The transcription factors responsible for the appearance of the exhausted phenotype in T cells have recently been identified and include NFAT, TOX, and NR4A family members^24–30^. Mechanistically, it was proposed that NFAT activation resulted from unbalanced Ca^2^ signalling due to defective co-stimulation^24^. NFAT then behaved as an initiating transcription factor by further driving the expression of TOX and NR4As^26,31^. By contrast, the mechanisms that deteriorate NK cell function during chronic infections and how they relate to the phenomenon of exhaustion as defined in the T cell field remain weakly defined.

To shed light on these issues, we established a cohort of CHB patients and healthy donors (HD) and validated the dysfunctional state and altered phenotype of circulating NK cells. Using flow cytometry, we showed that IL-15-mediated activation of the AKT/mTOR pathway was blunted in NK cells of CHB patients. However, this inadequacy did not translate into obvious metabolic defects. To identify the molecular mechanisms leading to dysfunction in an unbiased manner, we performed a transcriptome analysis of circulating NK cells of HD and CHB patients. We found that NK cells of CHB patients presented some of the key molecular hallmarks of exhausted T cells i.e. overexpression of transcription factors such as TOX or NR4A-family and their ICP targets. Furthermore, we uncovered a transcriptional signature implicating the activation of a partner-less NFAT, another characteristic of exhausted T cells. Mechanistically, this suggested that NK cells were submitted to unbalanced Ca^2+^ signalling in CHB patients. In order to test whether such unbalance could induce dysfunction, we induced Ca^2+^ flux in isolation in control NK cells. This treatment reproduced the dysfunction observed in CHB patients, thus providing molecular insight in the regulation of the dysfunctional state. Altogether, these data distinctly show that circulating NK cells in CHB patients exhibit key molecular features reminiscent of T cell exhaustion, a knowledge that could inform future immunotherapy strategies.

## Results

### NK cell functionality is impaired in CHB patients

We constituted a cohort of CHB patients and HD controls. Clinical parameters and statistics are presented in Table 1. The 32 patients constituting our cohort are in the immune inactive phase characterized by persistent HBV infection of the liver, absence of significant necroinflammatory disease (data not shown), low serum HBV DNA levels and normal serum aminotransferases. HD controls were sex and age matched to CHB patients. Similar to previously described cohorts, a higher proportion of CHB patients than HD were seropositive for HCMV (Table 1 and Supplemental Figure 1A), a characteristic that led to higher representation of the adaptive NKG2C^+^ subset in the NK cell population (Supplemental Figure 1B). Peripheral blood mononuclear cells (PBMCs) were isolated and NK cells identified as CD56^+^CD7^+^/CD3/19/14/4^-^ by flow-cytometry (Supplemental Figure 2). NK cell effector capacities were then analysed. Four different effector functions were measured: cytotoxicity and the production of three cytokines (IFN-γ, MIP1-β, and TNF-α). We measured these readouts in response to cell lines activating different receptors: K562 and Granta coated with Rituximab. As shown in Figure 1A, NK cells of CHB patients exhibited normal degranulation in response to K562 stimulation. Yet, basal levels of CD107a were increased in NK cells from CHB patients suggestive of recent stimulation. As CD107a exposure is only a surrogate marker of cytotoxicity, we directly measured cytotoxic capacity against K562 cells. For this purpose, we used a K562 sub-clone expressing the nano-luciferase enzyme, which, upon cell lysis is released in the culture medium allowing accurate quantification of cell death^32^. As shown in Figure 1B, the cytotoxic capacity of NK cells was not affected by HBV infection. In contrast, a significantly decreased capacity to produce IFN-γ and MIP1-β upon K562 stimulation was observed (Figure 1C). TNF-α secretion capacity was also decreased; however, this did not reach statistical significance. We then stimulated NK cells with Granta cells coated with Rituximab. To our knowledge this stimulus has not been previously tested in CHB patients. Degranulation was normal in CHB patients as measured by CD107a surface exposure (Supplemental Figure 3), while IFN-γ production was decreased (Figure 1D). Production of MIP1-β and TNF-α was comparatively less affected. In parallel, cytokine production was also measured in response to IL-12/18 stimulation (Figure 1E). No significant difference in IFN-γ, MIP1-β or TNF-α production was observed between CHB and HD. Overall, our results demonstrate a defect in cytokine secretion by NK cells from CHB patients when stimulated with MHC-I deficient or antibody-coated targets but not with IL-12/18.

**Table 1:**
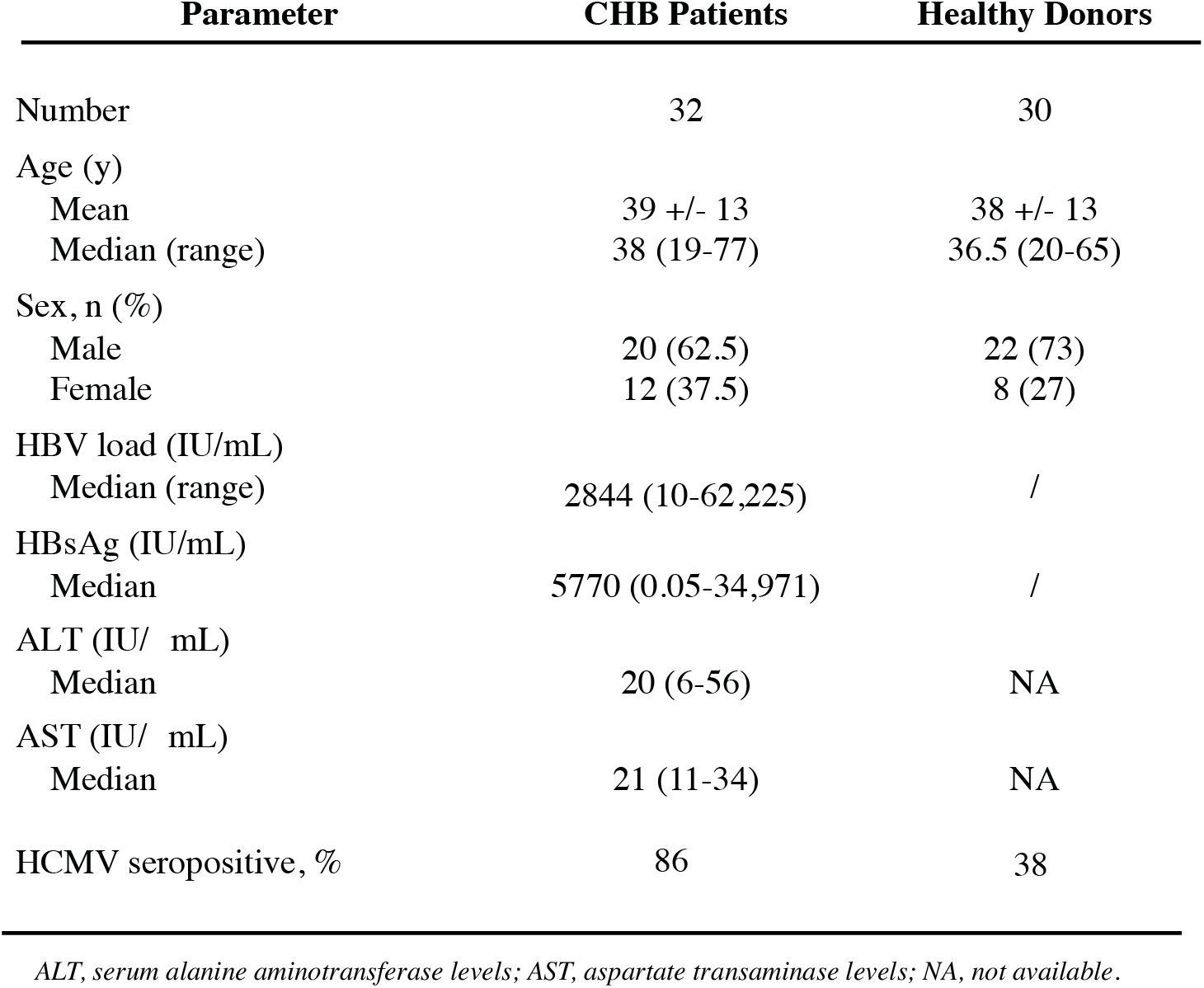
Characteristics of patients and HD enrolled in the study.

| Parameter | CHB Patients | Healthy Donors |
| --- | --- | --- |
| Number | 32 | 30 |
| Age (y) |  |  |
| Mean | 39 +/- 13 | 38 +/- 13 |
| Median (range) | 38 (19-77) | 36.5 (20-65) |
| Sex, n (%) |  |  |
| Male | 20 (62.5) | 22 (73) |
| Female | 12 (37.5) | 8 (27) |
| HBV load (IU/mL) |  |  |
| Median (range) | 2844 (10-62,225) | / |
| HBsAg (IU/mL) |  |  |
| Median | 5770 (0.05-34,971) | / |
| ALT (IU/ mL) |  |  |
| Median | 20 (6-56) | NA |
| AST (IU/ mL) |  |  |
| Median | 21 (11-34) | NA |
| HCMV seropositive, % | 86 | 38 |

ALT, serum alanine aminotransferase levels; AST, aspartate transaminase levels; NA, not available.

**Figure 1:**
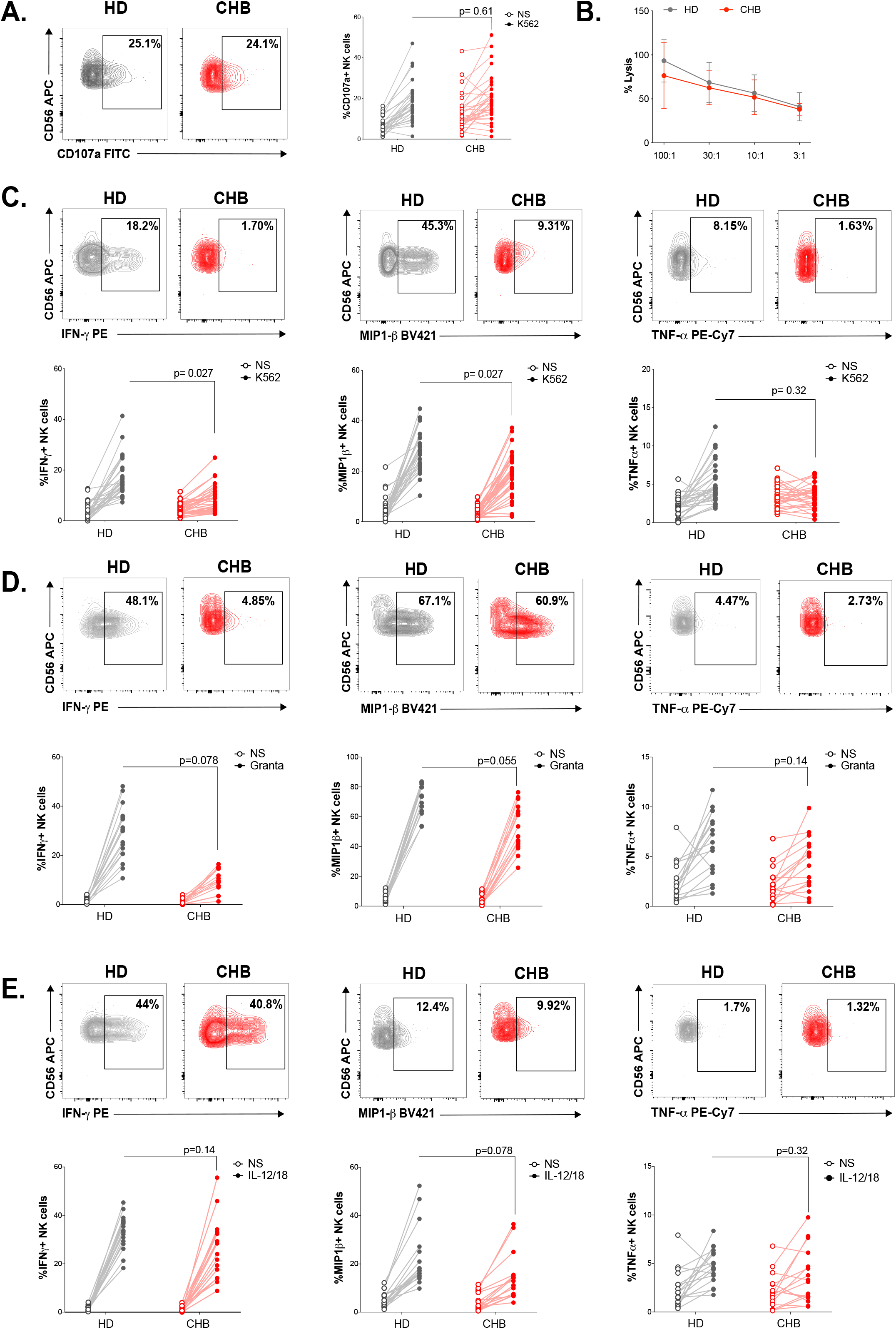
NK cell functionality is impaired in CHB patients. (A) PBMCs from HD (n=30) or CHB patients (n=32) were co-cultured with K562 during 4 hours and the proportion of NK cells expressing CD107a was determined by immunostaining. Representative flow-cytometry plots as well as proportion of CD107a+ NK cells is shown for each individual. (B) PBMCs from HD (n=17) or CHB patients (n=17) were cocultured with K562 Nanoluc+ at the indicated effectors:targets ratios during 4 hours. Supernatants were then collected to measure bioluminescence. Shown in the figure is the average bioluminescence +/-SD in an experiment with 5 HD and 5 CHB representative of four independent experiments. PBMCs from HD (n=30) or CHB patients (n=32) were co-cultured with K562 (C), Rituximab coated Granta (D) or with IL-12 and IL-18 (10 ng/mL each) during 4 hours. Intracellular stainings for the indicated cytokines were performed. Representative flow-cytometry plots as well as proportion of NK cells expressing the indicated molecule is shown for each individual (C= n=30 HD and n=32 CHB, D and E, n=17). Statistical analysis was performed by logistic regression as described in the Materials and Methods section and adjusted p-values are indicated on the graph. NS: non stimulated.

### NK cells from CHB patients display an altered phenotype

Next, we measured the percentage of total NK cells as well as the representation of subsets defined by CD56 expression levels following the gating strategy depicted in Supplementary Figure 2. We observed that the overall percentage of NK cells was decreased in CHB patients but this did not impact the relative representation of CD56^bright^ vs CD56^dim^ subsets (Figure 2A). In order to determine whether the poor functionality could be explained by altered expression of certain activating or inhibitory receptors, we characterized the phenotype of circulating NK cells in CHB patients compared to HD. The expression of activating receptors CD16 and NKp30 was significantly decreased while that of NKG2D was increased in CHB patients (Figure 2B); the expression of NKp46 and DNAM activating receptors was not different between HD and patients (Figure 2B). We also observed an overall reduction in the expression of the inhibitory receptors CD160, KLRG1 and NKG2A. However, this variation was statistically significant only for CD160 (Figure 2C). During NK cell education, the expression of self-engaged inhibitory receptors impacts the content of cytotoxic granules^33^. We thus measured the level of the cytolytic proteins Perforin and Granzyme B in NK cells of CHB patients and HD. We did not detect any difference in the expression of these molecules in accordance with the fact that cytotoxic capacities are preserved (Figure 2D). Collectively, our results show that dysfunctional NK cells in CHB patients have an altered expression of both activating and inhibitory NK cell receptors but a normal expression of cytotoxic molecules.

**Figure 2:**
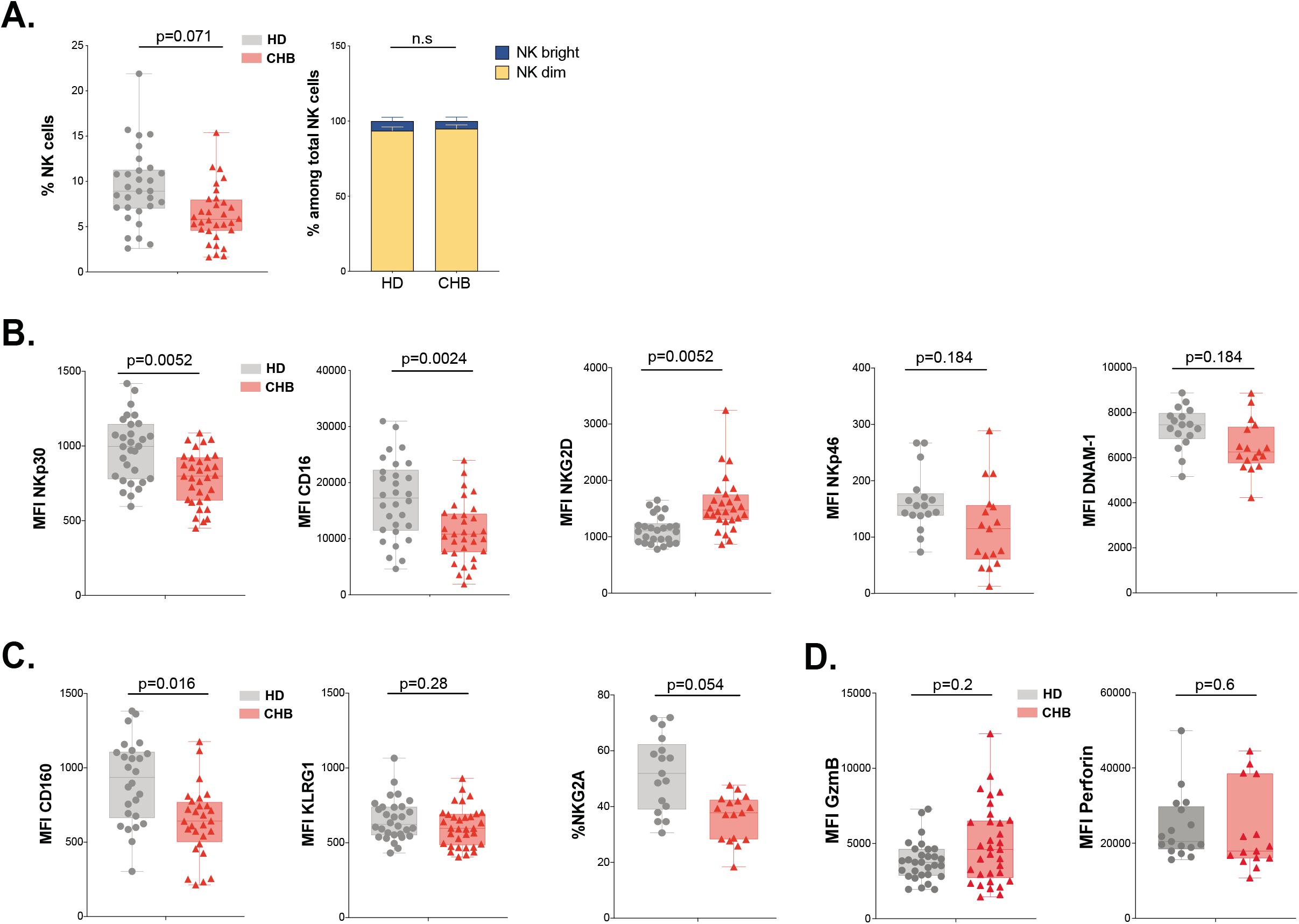
NK cell from CHB patients display an altered phenotype. (A) PBMCs from 30 HD and 32 CHB samples were stained with live/dead, CD4, CD14, CD19, CD3, CD7 and CD56 antibodies. The percentage of NK cells in each sample (+/- SD), and the proportion of CD56^bright^ versus CD56^dim^ NK cells (+/- SD), determined by flow cytometry, are shown. (B and C) Mean fluorescence intensity (MFI) of indicated NK activating (B) and inhibitory (C) receptors was determined by flow cytometry on total NK cells among PBMCs from HD or CHB patients. The mean expression (+/-SD) as well as values for each individual are represented, n=30 HD and 32 CHB samples for NKp30, CD16, NKG2D, KLRG1 and CD160, n=17 HD and 17 CHB samples for NKp46, DNAM-1 and NKG2A. (D) Mean fluorescence intensity (MFI) of Granzyme B and Perforin were determined by flow cytometry on total NK cells among PBMCs from HD or CHB patients. The mean expression (+/-SD) as well as values for each individual are represented, n=30 HD and 32 CHB samples for Granzyme B, n=17 HD and 17 CHB samples for Perforin. Adjusted p-values are indicated on the graph. n.s: non significant.

### mTOR activation is impaired in NK cells from patients

We have previously shown that NK cell responsiveness relies on the activity of the kinase mTOR^20^. In addition, the AKT/mTOR pathway is frequently blunted in exhausted T cells^23^. We thus measured mTOR activation in response to IL-15 stimulation in NK cells from CHB patients and HD. mTOR takes part in two distinct complexes: mTORC1 and mTORC2. In order to measure the activity of both complexes, we quantified the phosphorylation level of the ribosomal protein S6 (pS6) as well as the phosphorylation level of the kinase AKT on Ser473 (pAKT) downstream mTORC1 and mTORC2 respectively. In addition, we measured the phosphorylation of STAT5 (pSTAT5) as a control as it is induced by IL-15 stimulation while being independent of mTOR activation^18^. CD56^bright^ NK cells are highly sensitive to IL-15 treatment and strongly induce mTOR activity^34^, we thus focused our analysis on this population to maximise sensitivity. As depicted in Figure 3A, basal levels of pS6 were not altered in CHB patients compared to HD, yet upon IL-15 stimulation, mTOR activation was reduced in NK cells from CHB patients as shown by the decreased phosphorylation of both S6 and AKT. This defective induction was specific to the mTOR pathway as IL-15 triggering of pSTAT5 was not affected (Figure 3A). We reported that TGF-β is a potent negative regulator of the mTOR pathway in NK cells^19^ and reports indicate higher seric concentration of TGF-β in CHB patient^10,11^. We thus measured active TGF-β1 concentration in the serum of HD and CHB patients. Circulating TGF-β1 levels were indeed significantly higher in CHB patients (Figure 3B). However, no correlation was observed between TGF-β1 concentration and pS6 level neither at basal nor after IL-15 stimulation in the CHB patients group (data not shown). Higher TGF-β levels could thus participate in the reduction of mTOR activity in CHB patients but it is likely that other parameters take part in this phenomenon. As mTOR is a key regulator of metabolic networks, we investigated possible metabolic defects in NK cells of CHB patients. For this purpose, we first quantified cellular size and granularity, correlates of metabolic activity. As shown in Figure 3C, NK cell size was not affected in CHB patients (forward scatter (FSC) parameter) while they presented decreased granularity (side scatter (SSC) parameter). Furthermore, two nutrient transporters regulated by mTOR^18^, the heavy chain of system L amino-acid transporter (CD98) and the transferrin transporter (CD71), were also expressed at equivalent levels (Figure 3D). Previous reports have shown that alterations in mitochondrial activity were linked to lymphocytes exhaustion^35,36^. Hence, we quantified the global mitochondrial mass using the Mitotracker dye, and their production of reactive oxygen species (ROS) using the MitoSox dye, yet we did not observe any change between the two groups (Figure 3E). Overall these results demonstrate that circulating NK cells from CHB patients have a blunted capacity to activate the mTOR pathway that could potentially affect signalling through activating receptors. However, deregulation of mTOR did not translate into measurable basal metabolic changes.

**Figure 3:**
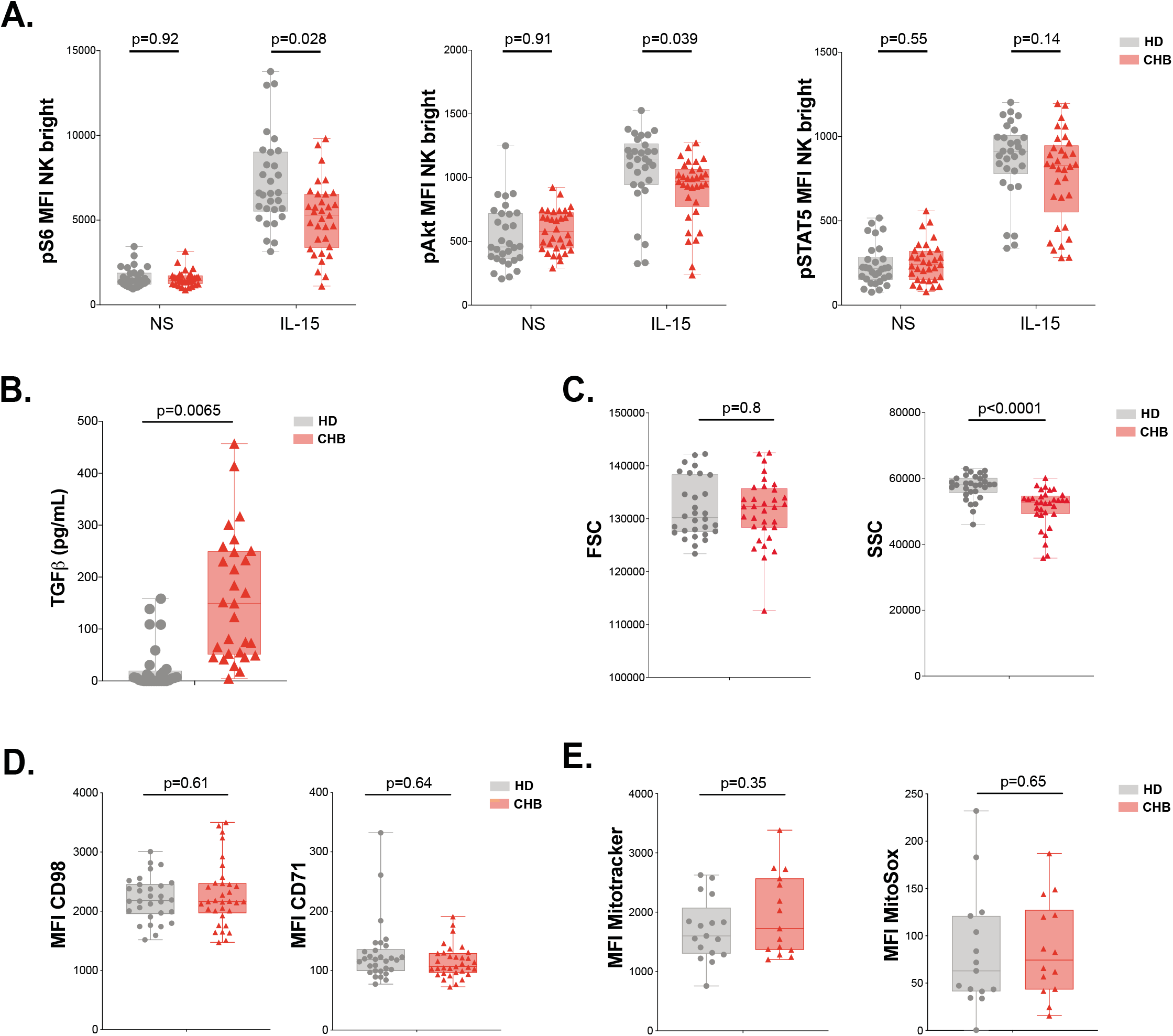
mTOR activation is impaired in NK cells from patients. (A) PBMCs from HD (n=30) or CHB patients (n=32) were stimulated with IL-15 at 100 ng/mL for 30 min. prior to phospho-epitope staining (pS6 Ser235/236, pAKT S473 and pSTAT5 Y694). The mean expression (+/-SD) as well as MFI values for each individual are represented, n=30 HD and 32 CHB (B) Active TGF-β1 levels were measured in serum samples from 30 HD and 32 CHB patients and the median (+/-SD) as well as values for each individual are represented. (C) FSC-A and SSC-A parameters on NK cells were measured by flow cytometry in 30 HD and 32 CHB samples. The mean of the MFI (+/-SD) as well as MFI values for each individual are represented. (D) MFI of CD98 and CD71 was determined by flow cytometry on total NK cells from HD or CHB patients. The mean expression (+/-SD) as well as values for each individual are represented, n=30 HD and 32 CHB samples. (E) PBMCs from 17 HD and 17 CHB patients were stained with MitoSOX Red and Mitotracker Green. Mean (+/-SD) as well as MFI values for each individual for the analysed marker are represented. Adjusted p-values are indicated on each graph.

### RNAseq analysis identifies an exhaustion-like signature in patient NK cells

We next took a non-hypothesis driven approach in order to seize the molecular mechanisms responsible for NK cell deregulation in CHB patients. For this purpose, we performed RNAseq on NK cells sorted from four CHB patients and five HD using the previously described gating strategy (Supplemental Figure 2). As shown in Figure 4A, PCA analysis of the results showed a clear separation between HD and CHB patients both on PC1 and 2, with 38% and 24% variance respectively, thus motivating further analysis of the results. Consistent with the fact that patients in this cohort are co-infected with HCMV, we found characteristics of adaptive NK cell populations in the differentially expressed genes (DEG) such as decreased *FCER1G, ZBTB16* (*PLZF*) or cytokine receptors mRNAs and increased *GZMH, KLRC4* or *CRTAM*^37^. In order to work with DEG that really reflected CHB impact, we filtered out genes that were significantly regulated in adaptive NK cells, as defined in a previous study^37^. This process identified 253 up-regulated and 163 down-regulated genes specific of HBV infection in CHB patients (FC>2 and adjusted p-value<0.05) (Figure 4B). We then analysed both gene lists using the online gene annotation tool Metascape^38^. No significant enrichment was found in the list of down-regulated genes. In contrast, analysis of the up-regulated genes retrieved Gene annotation terms that were consistent with ongoing viral infection such as “Viral life cycle” or “Hepatitis B” (Figure 4C, a complete version of the analysis is given in Supplemental Figure 4). Interestingly, some of the enriched terms referred to immune processes that are negatively impacted in NK cells of CHB patients such as “cytokine production”, “cytokine-mediated signalling”, “phosphorylation”, and “Protein kinase B (AKT) signalling”. We also noted that “T cell activation” was one of the enriched terms suggesting commonalities in the transcriptional regulation of NK and T cell responses. Moreover, we found that dysfunctional NK cells up-regulated several canonical genes of the T cell exhaustion-program, notably immune checkpoints or their ligands such as LAG3 and CD274 (PD-L1), or transcription factors such as EGR2 and 3, NR4A2 and TOX^25–29,39,40,31^ (Figure 4D). This observation prompted us to rigorously test whether the exhaustion transcriptional program was indeed undertaken by NK cells. To this aim, we performed Gene set enrichment analysis (GSEA) using two independent datasets defined in exhausted CD8 T cells in a context of chronic viral infection^41,42^. As depicted in Figure 4E, transcripts of these datasets were indeed strongly enriched in NK cells of CHB patients. This included TOX that we already identified among the genes significantly overexpressed in CHB patient NK cells (Figure 4D). This transcriptional regulator has recently been described as a key inducer of the exhausted gene signature allowing persistence of exhausted T cells^25–29^. We thus tested whether the TOX-induced gene signature was differentially expressed in HD vs CHB patients. We detected a significant enrichment of this signature in genes up-regulated in HBV patients (Figure 4F). In summary, NK cells of CHB patients display a transcriptional signature similar to the one observed in exhausted T cells induced by chronic viral infections. Furthermore, our data point to the involvement of the transcription factor TOX in driving NK cell dysfunction.

**Figure 4:**
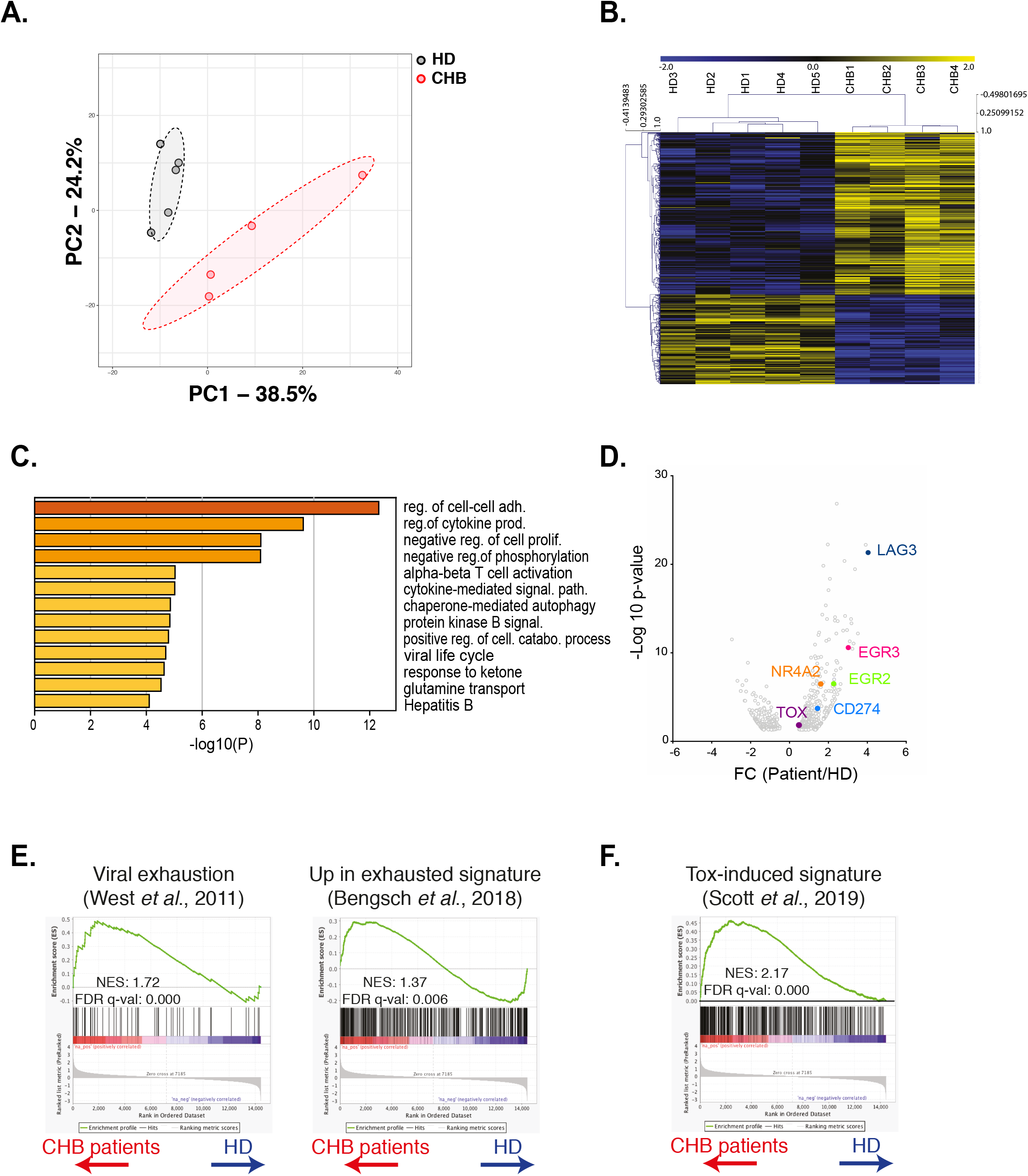
RNAseq analysis identifies an exhaustion-like signature in patient NK cells. (A) Principal Component Analysis of the RNAseq data is shown. (B) Heatmap of the DEG genes between HD and CHB. (C) Gene Ontology analysis of DEG up-regulated in CHB patients using Metascape. Selected terms are shown among the most significant ones. (D) Volcano Plots of the DEG highlighting genes belonging to the T cell exhaustion pathway. (E and F) GSEA plots comparing HD and CHB patients are shown for the indicated gene sets. The normalised enrichment scores (NES) and FDR q-values are indicated.

### Validation of the exhausted phenotype at the protein level

In order to validate the exhausted signature at the protein level, we recruited a second cohort constituted of 10 CHB patients and 9 HD independent from the first cohort. On this validation cohort, we first measured the intracellular expression of TOX in NK cells. Expression of transcription factors of the TOX family has previously been associated to early stages of NK cell development^43–46^, however, whether it is associated to the acquisition of a dysfunctional phenotype is unknown. As shown in Figure 5A, NK cells from CHB patients presented higher expression of TOX validating the RNAseq data. Increased TOX expression was seen mainly in the CD56^dim^ subset in CHB patients. In murine CD8 T cells, *Tox* invalidation abrogates the exhaustion program and in particular the expression of ICP such as LAG3, TIGIT, TIM3, 2B4, PD1 and CD39^25,27,28^. We observed that LAG3 was the most up-regulated gene in NK cells from CHB patients (Figure 4C). At the protein level, we also detected increased surface expression of LAG3 in NK cells from CHB patients compared to HD (Figure 5B). Of note, LAG3 and TOX protein levels were highly correlated in CHB patients specifically, further highlighting their functional link (Figure 5C, R^2^ = 0.88). Despite the fact that other ICP transcripts targeted by TOX were not significantly deregulated in our RNAseq analysis, we measured their protein expression. As shown in Figure 5D, TIGIT was up-regulated while TIM3 was down-regulated and 2B4 expression was unchanged in NK cells from CHB patients compared to controls. Some CHB patients presented limited but detectable PD1 expression above the average of HD. CD39 was not expressed (data not shown). In murine CD8 T cells, the transcription factor T-BET limits the expression of PD1^47^. In addition, viral induced CD8 T cell exhaustion has been linked to a decrease in T-BET and an increase in the expression of the closely-related T-box family transcription factor EOMES^48^, a point also validated in human during HIV-1 infection^49^. It is also reported that T-BET and EOMES are both required for NK cell differentiation and acquisition of effector functions^50^. We observed in CHB patients that T-BET was indeed significantly down-regulated while EOMES was unchanged (Figure 5E). Our data confirm the results obtained by RNAseq, demonstrating that NK cells from CHB patients display an exhaustion-associated signature at the protein level.

**Figure 5:**
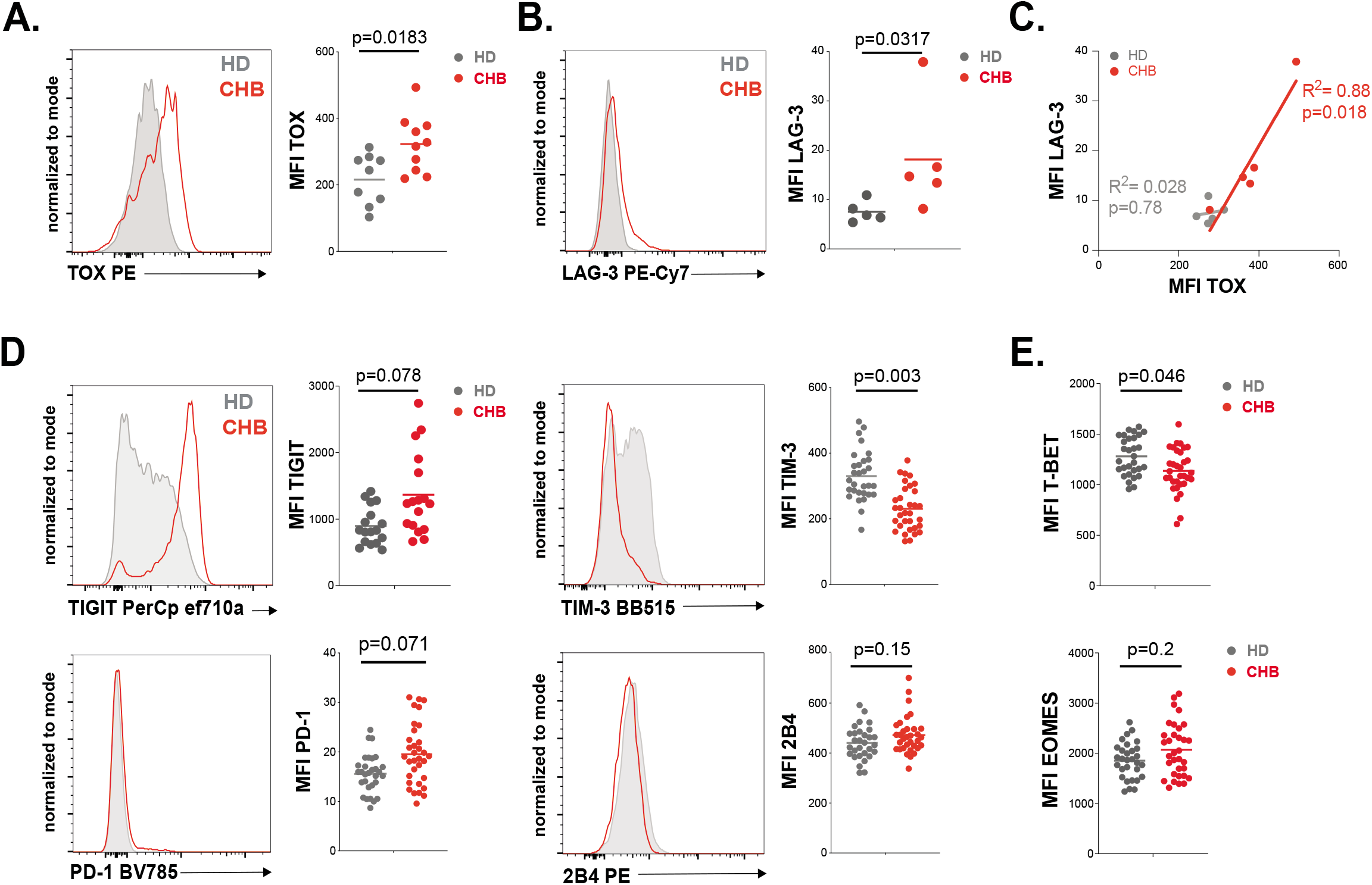
Validation of the exhausted phenotype at the protein level. (A) Intracellular staining for TOX was performed on PBMCs from 9 HD and 10 CHB samples and the MFI measured. A representative FACS histogram overlay as well as the average MFI and individual values for each sample are represented. (B) LAG-3 expression was measured by flow cytometry on PBMCs from 5 HD and 5 CHB samples. A representative FACS histogram overlay as well as the average MFI and individual values for each sample are represented. (C) Linear regression plots showing the correlation between TOX MFI and LAG-3 MFI using 5 HD and 5 CHB samples. The R^2^ and p-value calculated by linear regression are indicated. (D) ICP expression was measured by flow cytometry on NK cells from PBMCs of 17 HD and 17 CHB samples (TIGIT) or 30 HD and 32 CHB patients (TIM-3, PD-1 and 2B4). A representative FACS histogram overlay for each molecule as well as the average MFI and individual values for each sample are represented. (E) T-BET and EOMES expression were measured by intracellular staining on CD56^dim^ NK cells of 30 HD and 32 CHB samples. The average MFI and individual values for each sample are represented. Adjusted p-values are indicated on the graph.

### RNAseq and *in vitro* modelling indicate that NK cell dysfunction is due to unbalanced Ca^2+^ signalling

TOX controls the expression of phenotypic characteristics in exhausted T cells. It is however the effector of a more global transcriptional program. The transcription factors initiating this program appear to be NFAT-family members^24,51^. Indeed, it has been proposed that chronic TCR activation leads to unbalanced Ca^2+^-related signalling and consequently nuclear translocation of an excess of NFAT relative to its AP-1 partners. This partnerless NFAT binds to and transactivates a specific subset of genes, distinct from the canonical subset regulated by NFAT:AP1 heterodimers and substantially overlapping the transcriptional program of exhausted cells^24^. In particular, transcription factors such as *TOX* or the *NR4A* family are validated targets of partnerless NFAT^26^. Based on this knowledge, we performed GSEA analysis using a gene set previously defined by Martinez *et al*., to be regulated by partnerless NFAT in context of altered Ca^2+^ signalling^24^. We found that this gene set was enriched in NK cells of CHB patients (Figure 6A). This suggested that, in CHB patients, NK cells are subjected to an 2+ 2+ unbalanced Ca^2+^ signalling. To functionally test whether an altered Ca^2+^ signalling would endow control NK cells with characteristics of CHB patient NK cells, we pre-treated PBMCs from HD for 16h with different concentrations of ionomycin, a Ca^2+^ ionophore. We then measured NK cell effector functions: degranulation and cytokine expression in response to a K562 challenge. As previously published on NK cell lines^52^, a dose of 100nM ionomycin rendered primary NK cells unable to perform any of the tested functions (Figure 6B). In contrast, at 1nM, no detectable effect on degranulation was observed, yet, it was sufficient to impair IFN-γ production, thus reproducing the functional dichotomy observed in CHB patients. The production of MIP1-β and TNF-α showed intermediate behaviours in accordance with CHB patients’ phenotype. This was achieved without negative impact on viability (data not shown). Overall, these results show that stimulation of Ca^2+^ flux in isolation can reproduce the dysfunctional phenotype of NK cells observed in a context of CHB infection. These data, in association with transcriptional enrichment of partnerless NFAT-dependent transcripts, strongly suggest that NK cell dysfunction observed in CHB patients is the result of unbalanced Ca^2+^ signalling. This further indicates that common molecular mechanisms govern T cell exhaustion and NK cell dysfunction in contexts of chronic stimulation.

**Figure 6:**
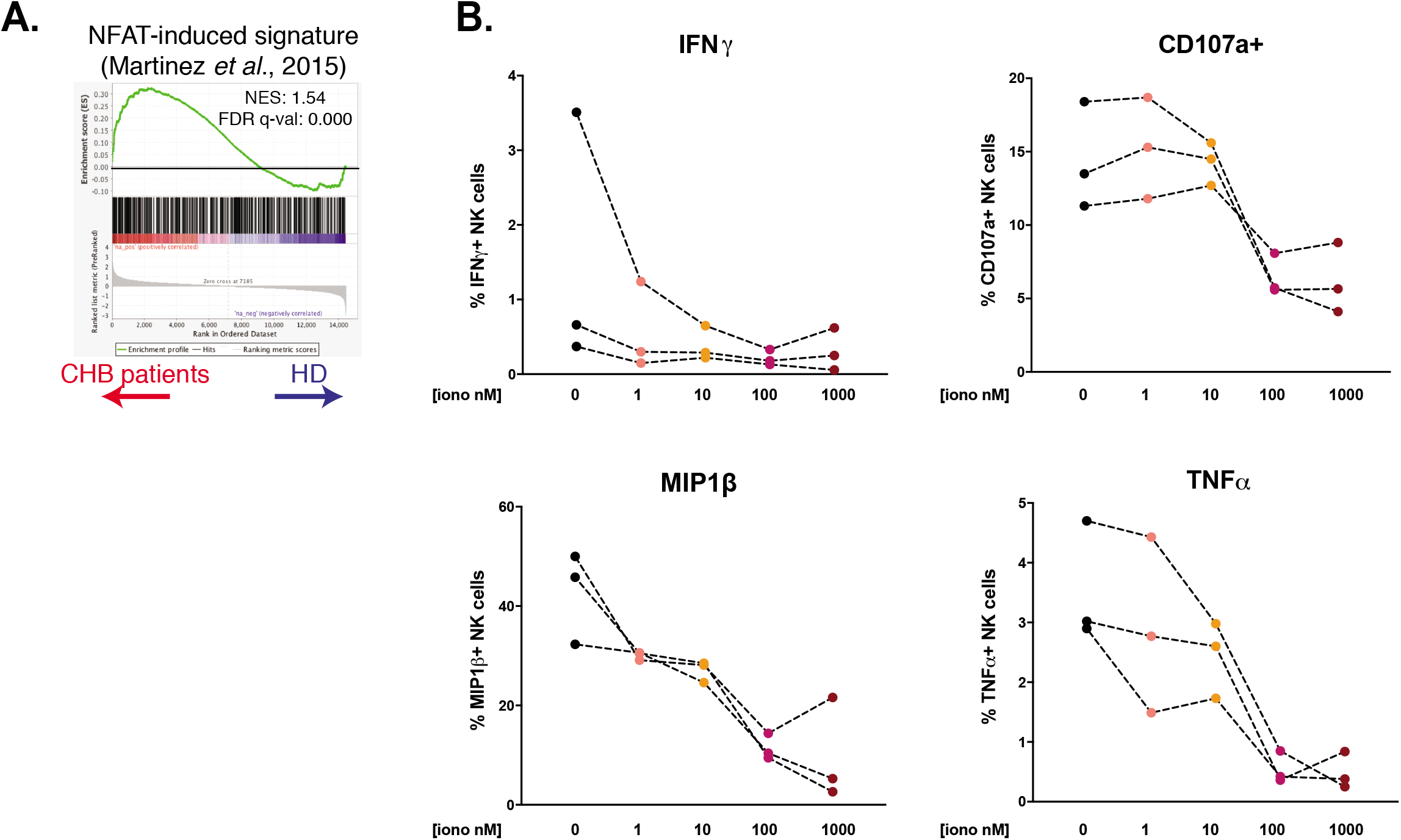
RNAseq and *in vitro* modelling suggest that NK cell dysfunction is due to unbalanced Ca^2+^ signalling. (A) A GSEA plot comparing HD and CHB patients is shown for the gene set regulated by partnerless NFAT. The normalised enrichment score (NES) and FDR q-values are indicated. (B) PBMCs from 3 HD were incubated with ionomycin at the indicated concentration for 16 hours. NK cell capacity to degranulate or produce the indicated cytokines upon K562 stimulation was then measured by flow cytometry. The average (+/-SD) and individual values of the proportion of positive NK cells is shown. The experiment has been performed twice.

## Discussion

In the present study, we explored the impact of CHB on peripheral NK cell function and phenotype and uncovered a profound convergence in the transcriptional mechanisms governing T cell exhaustion and NK cell dysfunction.

We first confirmed previous reports showing that NK cell production of cytokines, chief among them IFN-γ, is blunted in response to target cell stimulation, while cytotoxic functions are not affected in the same settings^9–12^. This functional dichotomy could be particularly relevant since an early study of acute HBV infection in chimpanzee suggested that viral clearance was mediated by non-cytopathic antiviral effects^53^. We can thus hypothesize that loss of cytokine-mediated control contributes to HBV escape and establishment of chronicity.

NK cells of CHB patients constituting our cohort respond normally to IL-12/18 challenge in accordance with previous findings^11^. This is however in contrast with other reports^10,12^. This discrepancy could stem from differences in the abundance of HCMV-induced adaptive NK cells in the respective cohorts. Indeed, these cells present impaired responses to IL-12/18^37^ and, CHB patients being frequently co-infected with HCMV^54^, NKG2C^+^ adaptive NK cells are significantly more represented in CHB patients^55,56^. In our cohort, we observed that patients presenting the highest NKG2C^+^ frequency were less responsive to IL-12/18 (data not shown). Interestingly, the overall functionality of HCMV-induced adaptive NK cell is not affected by HBV^56^. It thus seems that both viruses engage non-overlapping NK cell subsets.

We report here that the functional impairment imposed by CHB significantly imprints the NK cell transcriptome. A recent study was performed on HBV-infected patients from all four phases of infection^57^. Very few transcripts were significantly deregulated in the inactive carriers compared to HD controls. In particular, no IFN-I stimulated genes were upregulated, a finding we confirm with our dataset. Further mining our data, we found that the transcriptome of NK cells in CHB patients contained gene signatures defined in exhausted CD8 T cells arising in chronic viral infection models. This strongly indicates that a mechanism akin to exhaustion is at play to explain the dysfunctional phenotype of NK cells in CHB patients. In particular, we found that the gene signature defined by transcripts regulated by the transcription factor TOX were enriched in CHB patients. This transcription factor has recently been involved in the establishment of the exhausted phenotype in CD8 T cells, its expression being driven by chronic TCR stimulation^25–29^. TOX family has been associated to NK cell development^43–46^, however, to our knowledge, this is the first indication that TOX participates in NK cell dysfunction. These results further underscore the ontogenic and functional proximity of NK cells and CD8 T cells. Monitoring reversal of NK cell dysfunction in therapeutic settings aimed at reversing T cell exhaustion in chronic diseases should thus be considered. Mechanistically, it further suggests that direct chronic stimulation, perhaps through NK activating receptors, is responsible for the dysfunction. Reinforcing this hypothesis is the fact that we and others observed decreased NKp30 and CD16 expression on CHB patient NK cells suggesting chronic engagement^10,12^. The nature of the exact stimulus and receptor involved remains to be deciphered. Since we studied peripheral blood NK cells, a population largely distinct from liver resident NK cells^58^, it is unlikely that dysfunction results from direct stimulation by infected cells. Circulating HBs antigen is however present in complex with neutralizing antibodies^59^. These immune complexes could chronically stimulate NK cells through CD16 cross-linking. In this respect, it has recently been reported that influenza vaccination increases NK cell function inducing memory-like differentiation^60^. This property is based on the sensing of immune complexes by CD16^61^. Based on the CD8 T cell system where a given stimulus can give rise to functional memory cells in acute settings and to exhausted cells if it becomes chronic^62,63^, we can hypothesize that chronic stimulation of NK cells through CD16 progressively drives the exhausted signature seen in our RNAseq.

Given the molecular proximity we describe, we could expect exhausted NK cells to display other characteristics of exhaustion observed in T cells. A key feature when T cells progress to exhaustion is a defective metabolism and blunted activation of the mTOR pathway, a major coordinator of anabolic and catabolic pathways. In CHB patients in response to IL-15, we indeed observed a decrease in the activation of the mTOR pathway, quantified by lower mTORC1 and 2 activities. Importantly, this defect selectively affected the mTOR pathway since phosphorylation of STAT5, another signalling event downstream IL-15 Receptor (IL-15R), was normal. Such a depression in mTOR activity due to chronic activation is reminiscent of the situation that prevails in so called “non-educated” or “disarmed” hyporesponsive NK cells^20^. Of note, this similarity also points toward excessive stimulation of activating receptors, the mechanism envisaged to explain NK cell disarming^33,64,65^. How chronic stimulation leads to decreased mTOR activity remains to be investigated. Weaker mTORC1 activity could lead to impaired cell metabolism. However, we did not detect any significant metabolic defect in CHB patients using several readouts, in contrast to what has been reported for CD8 T cells in CHB patients^35^. This finding, associated to the fact that the cytotoxic capacity is spared and that expression of ICPs is low in most patients suggests either that exhaustion is at an early stage or that, unlike T cells, NK cells have a limited capacity to engage a full exhaustion program.

At the molecular level, our results point to a major role of Ca^2+^ and downstream NFAT signalling in the induction of NK cell dysfunction. Indeed, we detected a transcriptional signature indicative of improper activation of NFAT transcription factors in dysfunctional NK cells from CHB patients. In line with our findings, we devised an *in vitro* model based on unbalanced NK cell activation by activation of the Ca^2+^ pathway alone. Keeping in mind that exhaustion is only partial in CHB patients, we titrated ionomycin down to 1nM. At this dose, the only affected function in response to K562 stimulation was the ability to produce IFN-γ. Thus validating that this model phenocopies the functional dichotomy seen in circulating NK cells of CHB patients. Of note, such stimulus was also shown to induce hypo-responsiveness in T cells and NK cell lines reinforcing the parallel between both cell types^52,66,67^. Interestingly, the effect of ionomycin treatment can be adjusted so that higher concentration leads to more pronounced defects until loss of degranulation capacity.

How an imbalance in Ca^2+^ responses is triggered in CHB context remains an open question. In the T cell field, such an imbalance is proposed to be the result of defective co-stimulation^24^. However, in NK cells, the conceptual framework of stimulatory vs costimulatory receptors is not as clearly established. We hypothesize that a mechanism similar to a recently proposed model of disarming could be at play^33^. In this model continuous leakage of cytotoxic granules in response to chronic activating receptor triggering is involved. Since cytotoxic granules are part of the acidic Ca^2+^ stores, we can hypothesize that low-grade degranulation would lead to increased concentration of intracellular free Ca^2+^ and activation of NFAT. However, in contrast to CHB, disarming does not imprint the transcriptome of circulating NK cells^33,68^. It is thus likely that a combination of signals such as stimulation of activating receptors and increased levels of anti-inflammatory cytokines (TGF-β and IL-10) (this study and others^10,11^) shapes the NK cell phenotype in CHB patients.

In addition to their interest at the basic level, the findings we present here will aid to rationally design successful NK cell reinvigoration strategies that could contribute to viral elimination. Based on our findings, we would hypothesize that targeting the Ca^2+^ pathway or mechanistically linked molecules to re-establish a correct signalling balance could have a positive impact on effector functions. In this respect, a recent screen identified ingenol mebutate (IngMb), an activator of PKCs, for its ability to revert T cell exhaustion^69^. Mechanistically, this compound was able to complement ionomycin signal for efficient reinvigoration of virus-specific T cells. Indeed, PKCs are activated by a signalling branch parallel to Ca^2+^ signalling and participate in the activation of the transcription factor AP1, the missing partner of NFAT in exhausted cells^70^. PKCs activation could thus compensate the observed signalling defect and constitute a useful target to restore dysfunctional NK cells activity. More studies will be required to address this point.

In summary, we provide evidence that dysfunctional NK cells of CHB patients present a molecular signature similar to the one of exhausted T cells at the transcriptional and protein level. This signature is indicative of a signalling imbalance involving calcium. This could open the way toward original therapeutic targets.

## Materials and Methods

### Patients and Healthy donors

Peripheral blood samples from healthy subjects were obtained from the French blood agency (Etablissement Français du sang, Lyon, France). PBMCs from CHB patients and clinical assessments were obtained during routine hepatitis consultations. All participants provided written informed consent in accordance with the procedure approved by the local ethics committee (Comité de Protection des Personnes, Centre Hospitalier Universitaire de Limoges, Limoges, France) and the Interventional research protocol involving human samples (Code promotor LiNKeB project: 87RI18-0021). Patients and healthy donor characteristics are detailed in Table 1. All patients were diagnosed as inactive carriers according to the American Association for the Study of Liver Diseases (AASLD) guidelines for treatment of chronic hepatitis B^71^. HCMV seropositivity status of patients and healthy donors were determined by ELISA (Hôpital de la Croix-Rousse, Lyon, France).

### PBMCs isolation

Human PBMCs were separated from peripheral blood by Ficoll gradient centrifugation (Eurobio Laboratoires et AbCys) at room temperature (RT). Cells were then resuspended in heat-inactivated FCS with 20% DMSO, progressively cooled down to −80°C and stored in cryotubes in liquid nitrogen.

### Cell culture and treatments

Cells were cultured in RPMI 1640 medium (Invitrogen Life Technologies) supplemented with 10% of FCS, 2mM L-glutamine, 10 mM of penicillin/ streptomycin (HCL Technologies), and 1 mM sodium pyruvate (PAA Laboratories) and 20 mM HEPES (Gibco). For phosphorylated protein analysis, cells were stimulated with IL-15 (Peprotech, 100ng/mL) during 30 minutes at 37°C. In some experiments, cells were treated with increasing doses of ionomycin (Sigma) during 16 hours.

### Flow cytometry analysis

PBMCs were rapidly thawed in medium heated to 37°C and kept overnight at 4°C. Cells were then immunostained during 30 min at 4°C with the appropriate monoclonal antibodies detailed in Table S1. Intracellular staining of transcription factors and cytotoxic molecules was performed with Foxp3 Fixation/Permeabilization concentrate and diluent (eBioscience). Intracellular staining of cytokines and chemokines was performed with Cytofix/Cytoperm (BD Biosciences). Intracellular staining of phosphorylated proteins was performed with Lyse/Fix and Perm III buffers (BD Biosciences). Phosphorylated proteins were then stained during 40 minutes at RT. Flow cytometric analysis was performed on LSR Fortessa 5L (Becton-Dickinson). Fluorescence Minus One (FMO) controls were used to set the gates and data were analysed with FlowJo 10.5.0 software (Tree Star). Gating strategy is presented is Figure S2.

### NK cell stimulation

Human PBMCs were stimulated with recombinant human IL-12 (Peprotech) and IL-18 (R&D Systems) at 10 ng/mL each or co-cultured for 4 hours with K562 cells or with Granta cells coated with Rituximab at a 1.1 ratio in the presence of Golgi Stop (BD Biosciences). The percentage of NK cells positive for CD107a, MIP1-β, IFN-γ and TNF-α was then determined by flow cytometry.

### Cytotoxic assay

Human PBMCs were rapidly thawed in medium heated to 37°C and kept overnight at 4°C. Cells were then co-cultured for 4 hours at different Effectors:Targets ratios with K562 NanoLuc^+32^. Supernatant was collected and bioluminescence was measured using TECAN Instrument after addition of Furimazin, the Nanoluciferase substrate (Promega).

### Mitochondria analysis

Human PBMCs were incubated with MitoSOX Red (5μM) and Mitotracker Green (1 μM) (both from Molecular Probes, Life Technologies) in PBS during 10 minutes at 37°C before flow cytometry extra-cellular staining.

### Serum TGF-*β1* quantification

Active TGF-β1 serum levels in patients and healthy donors were measured using LEGEND MAX™ Free Active TGF-β1 ELISA Kit with Pre-coated Plates (Biolegend). The assay was run according to the manufacturer’s recommendations.

### RNA Sequencing

NK cells from 5 HD and 4 CHB were sorted as live/dead^-^/CD4^-^/14^-^/19^-^/CD3^-^/CD56^+^ cells by flow cytometry. Samples from HD were sorted in a BSL2 cytometry platform, (Anira cytométrie, SFR Biosciences, Lyon, France), whereas samples from CHB were sorted in a BSL3 cytometry platform (Toulouse, France). NK cells were then lysed in Direct-Zol (Ozyme). The RNA libraries were prepared according to the protocol of Picelli *et al*.^72^. Total RNA was purified using the Direct-Zol RNA Microprep Kit (Ozyme) according to the recommendations provided and was quantified using the QuantiFluor RNA system (Promega). One μl of 10 μM of oligo-dT primer and 1 μl of 10 μM of dNTPs were added to 0.3 ng of total RNA in a final volume of 2.3 μl. The Oligo-dTs were hybridized for 3 min at 72 ° C and a reverse transcription reaction was carried out as described in the Nature protocols. The complementary DNAs (cDNAs) were purified on AmpureXP beads (Beckman Coulter) and the quality was checked on a D5000 screening strip and analysed on a 4200 strip station (Agilent). Three ng of cDNA were labelled using the Nextera XT DNA sample preparation kit (Illumina). The labelled fragments were then amplified following PCR cycles and purified on AmpureXP beads (Beckman Coulter). The quality of the bank was checked on a D1000 screen tape and analysed on a 4200 tape station (Agilent). The sequencing of the banks was carried out by the GenomEast platform, member of the “France Génomique” consortium (ANR-10-INBS-0009).

### In silico analyses

PCA was performed using the R software (version 3.6.1) after data normalization and graphed with the ggplot2 package. We then obtained a first DEG list using an adjusted p-value <0.05 as a cutoff (750 genes). In order to keep DEG deregulated only by HBV infection in our subsequent analysis, we removed the ones known to be affected by HCMV infection. To identify genes expressed by NK cells and affected by HCMV infection, we used a previously published microarray dataset comparing conventional and adaptive NK cells^37^. In this HCMV dataset, DEG were identified as genes presenting a p-value below 0.005 (1357 genes in total). Elements common to the two DEG lists and that did not satisfy to the relation: *abs(logFC(HBV))>2*abs(logFC(HCMV))* were subtracted from the list of DEG obtained in our RNAseq study. We further identified genes showing a *FC(HBV)>2* to obtain the final list of DEG genes. This DEG list was used for heatmap and Metascape analysis. The heatmap was constructed using Multiple Experiment Viewer with Row Median centering of the data^73^. Functional annotations of DEG were performed with Metascape^38^ using default parameters. Regarding GSEA analysis, indicated gene sets of publicly available expression data were obtained. To statistically test whether these gene sets were enriched in specific conditions, we performed pairwise comparisons between HD and CHB patients’ conditions using the GSEA method (http://www.broad.mit.edu/gsea).

### Statistical analysis

Clinical data were processed with the R statistical environment. After cleaning empirically data from outliers, we transformed to Log scale the parameters that were showing log-normal distribution and kept others unchanged. Then we used a generalized linear model with binomial family (logistic regression model) in simple way, *i.e* one regressor at time, to quantify for each biological parameter independently the probability to be linked to our outcome variable (= to be HD or Patients). The results for all parameters were taken together to correct for the multiple testing using the Benjamini-Hochberg method. Graphical representations were done using Prism 5 (Graph-Pad Software) unless otherwise stated.

## Acknowledgments

We thank the SFR Biosciences (UMS3444/CNRS, ENSL, UCBL, US8/INSERM) facilities, in particular the Plateau de Biologie Expérimentale de la Souris, and the flow cytometry facility. We would like to thank Dr P.O Vidalain who provided the K562 NanoLuc, and Dr. Christophe Ramière for the determination of HCMV status in Hôpital Croix-Rousse. We also acknowledge the contribution of the GenomEast platform from Strasbourg, France. TW lab is supported by the Agence Nationale de la Recherche (ANR JC *SPHINKS* to TW and ANR JC *BaNK* to AM), the ARC foundation (équipe labellisée), the European Research council (ERC-Stg 281025), and receives institutional grants from the Institut National de la Santé et de la Recherche Médicale (INSERM), Centre National de la Recherche Scientifique (CNRS), Université Claude Bernard Lyon1 and ENS de Lyon. MM is the recipient of a fellowship from La Ligue Nationale contre le Cancer. This work was supported by l’Agence Nationale de Recherche sur les SIDA et les Hépatites virales (ANRS projects, ECTZ22398, ECTZ11169 and ECTZ19856 to UH) and Comité du Rhône LNCC (to UH).

## Author contributions

MM, MV, IT, and LB performed and analyzed experiments. MP, YR, MA, and GR assisted with experiments and SV supervised some experiments. VL and SA provided the patient cohorts. MG prepared CHB patient samples. UH, AM and TW conceived the study, designed and interpreted experiments. OA performed the biostatistical analysis. DD provided scientific advice and helped to design experiments. MM and AM wrote the manuscript with inputs from UH, and TW. All authors critically read the manuscript.

## Materials and Correspondence

Correspondence and material requests should be addressed to Thierry Walzer, Uzma Hasan or Antoine Marçais.

