## Supplemental figures and table for "Peripheral Natural Killer cells from chronic hepatitis B patients display molecular hallmarks of T cell exhaustion"

### Supplementary Figures and Table

**<u>Figure S1: HCMV status</u>**

(A) The concentrations of IgG anti CMV antibodies were measured in serum from patients and HD to determine HCMV status. (B) The percentage of NKG2C+ adaptive NK cells in patients and HD was determined by flow cytometry and represented according to HCMV status.

**<u>Figure S2:</u> Gating Strategy**

Gating strategy used for flow cytometry analysis: First, lymphocytes through SSC and FSC analysis were selected. Then doublets were eliminated as well as dead cells. We then removed $CD4^+$, $CD14^+$ and $CD19^+$ cells prior to gate on NK cells by CD56+ and CD3- cells. CD7 marker was used to confirm the gating and NK bright and dim were then separated.

**<u>Figure S3:</u> Metascape analysis of CHB patients up-regulated genes**

Full results for the Gene Ontology analysis of DEG up-regulated in CHB patients performed with Metascape and presented in Figure 4C.

**<u>Table S1:</u> Antibodies used**

**Figure S1.**

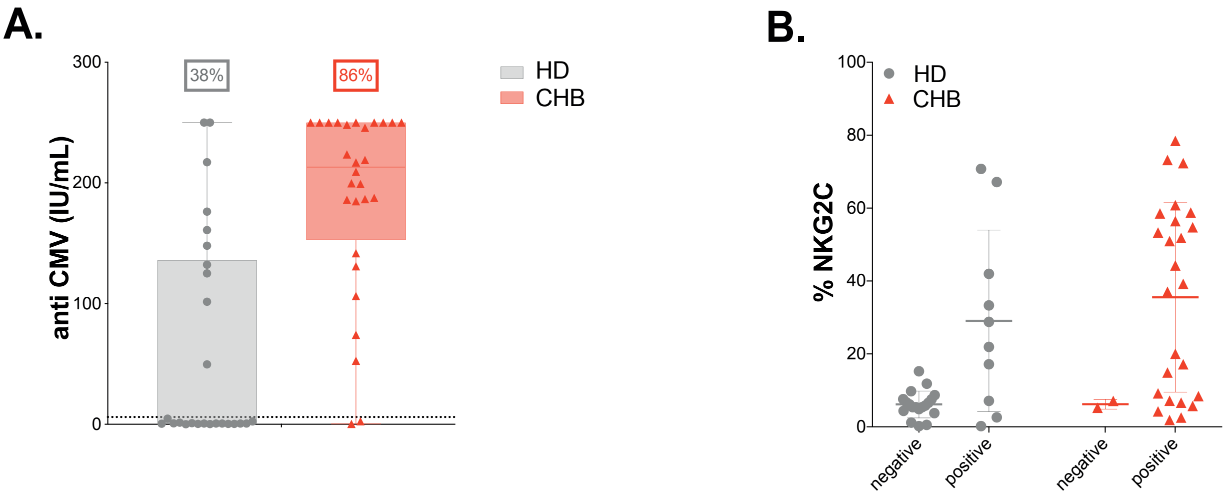


**Figure S2.**

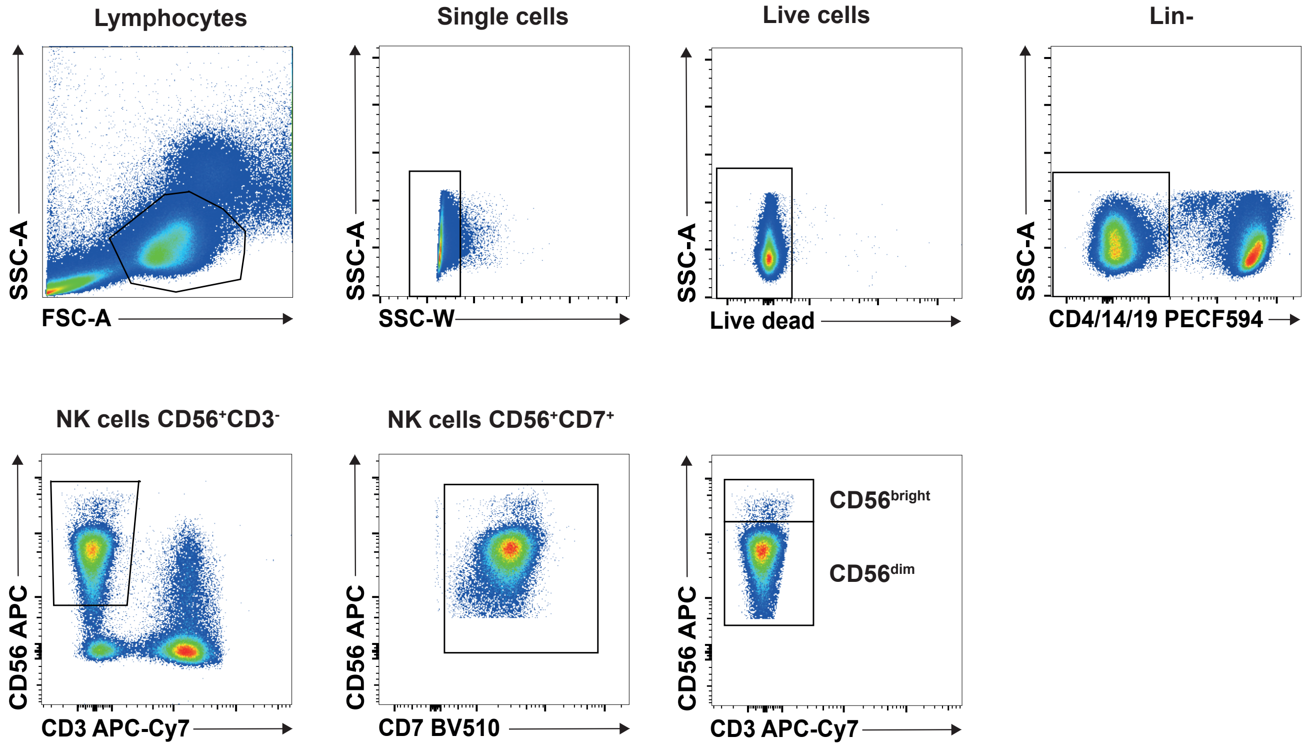


**Figure S3.**

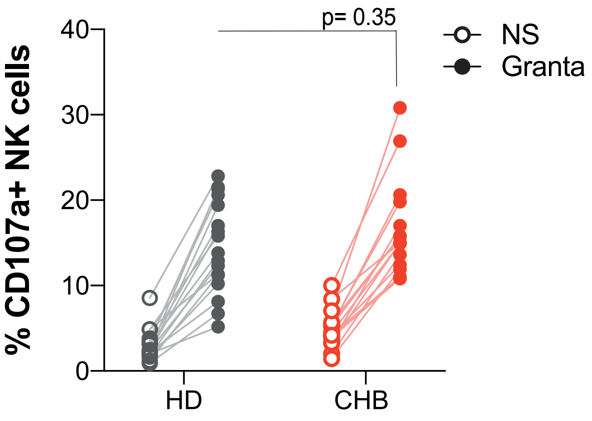

**Figure S4.**

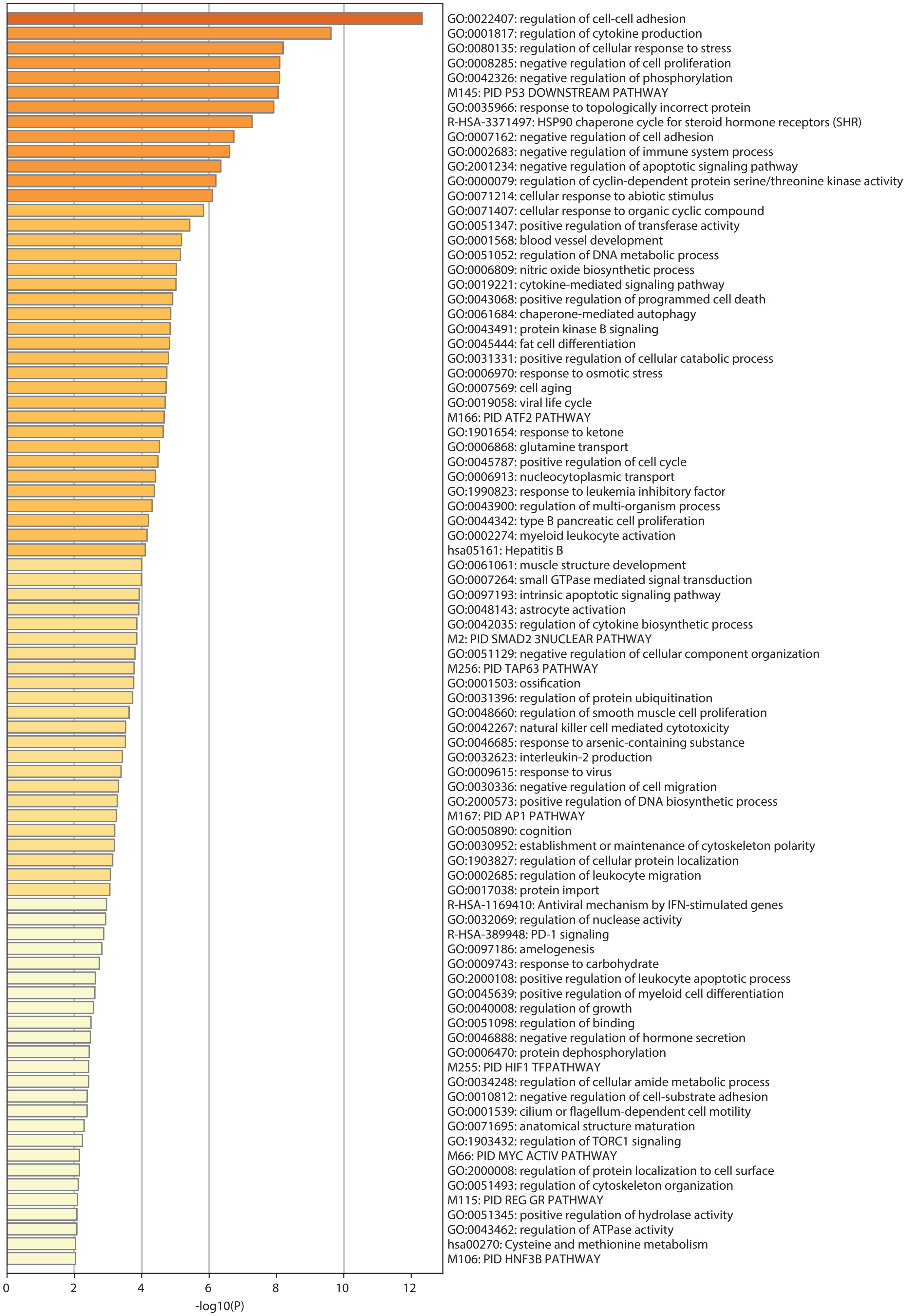
GO:0022407: regulation of cell-cell adhesion
GO:0001817: regulation of cytokine production
GO:0080135: regulation of cellular response to stress
GO:0008285: negative regulation of cell proliferation
GO:0042326: negative regulation of phosphorylation
M145: PID P53 DOWNSTREAM PATHWAY
GO:0035966: response to topologically incorrect protein
R-HSA-3371497: HSP90 chaperone cycle for steroid hormone receptors (SHR)
GO:0007162: negative regulation of cell adhesion
GO:0002683: negative regulation of immune system process
GO:2001234: negative regulation of apoptotic signaling pathway
GO:0000079: regulation of cyclin-dependent protein serine/threonine kinase activity
GO:0071214: cellular response to abiotic stimulus
GO:0071407: cellular response to organic cyclic compound
GO:0051347: positive regulation of transferase activity
GO:0001568: blood vessel development
GO:0051052: regulation of DNA metabolic process
GO:0006809: nitric oxide biosynthetic process
GO:0019221: cytokine-mediated signaling pathway
GO:0043068: positive regulation of programmed cell death
GO:0061684: chaperone-mediated autophagy
GO:0043491: protein kinase B signaling
GO:0045444: fat cell differentiation
GO:0031331: positive regulation of cellular catabolic process
GO:0006970: response to osmotic stress
GO:0007569: cell aging
GO:0019058: viral life cycle
M166: PID ATF2 PATHWAY
GO:1901654: response to ketone
GO:0006868: glutamine transport
GO:0045787: positive regulation of cell cycle
GO:0006913: nucleocytoplasmic transport
GO:1990823: response to leukemia inhibitory factor
GO:0043900: regulation of multi-organism process
GO:0044342: type B pancreatic cell proliferation
GO:0002274: myeloid leukocyte activation
hsa05161: Hepatitis B
GO:0061061: muscle structure development
GO:0007264: small GTPase mediated signal transduction
GO:0097193: intrinsic apoptotic signaling pathway
GO:0048143: astrocyte activation
GO:0042035: regulation of cytokine biosynthetic process
M2: PID SMAD2 3NUCLEAR PATHWAY
GO:0051129: negative regulation of cellular component organization
M256: PID TAP63 PATHWAY
GO:0001503: ossification
GO:0031396: regulation of protein ubiquitination
GO:0048660: regulation of smooth muscle cell proliferation
GO:0042267: natural killer cell mediated cytotoxicity
GO:0046685: response to arsenic-containing substance
GO:0032623: interleukin-2 production
GO:0009615: response to virus
GO:0030336: negative regulation of cell migration
GO:2000573: positive regulation of DNA biosynthetic process
M167: PID AP1 PATHWAY
GO:0050890: cognition
GO:0030952: establishment or maintenance of cytoskeleton polarity
GO:1903827: regulation of cellular protein localization
GO:0002685: regulation of leukocyte migration
GO:0017038: protein import
R-HSA-1169410: Antiviral mechanism by IFN-stimulated genes
GO:0032069: regulation of nuclease activity
R-HSA-389948: PD-1 signaling
GO:0097186: amelogenesis
GO:0009743: response to carbohydrate
GO:2000108: positive regulation of leukocyte apoptotic process
GO:0045639: positive regulation of myeloid cell differentiation
GO:0040008: regulation of growth
GO:0051098: regulation of binding
GO:0046888: negative regulation of hormone secretion
GO:0006470: protein dephosphorylation
M255: PID HIF1 TFPATHWAY
GO:0034248: regulation of cellular amide metabolic process
GO:0010812: negative regulation of cell-substrate adhesion
GO:0001539: cilium or flagellum-dependent cell motility
GO:0071695: anatomical structure maturation
GO:1903432: regulation of TORC1 signaling
M66: PID MYC ACTIV PATHWAY
GO:2000008: regulation of protein localization to cell surface
GO:0051493: regulation of cytoskeleton organization
M115: PID REG GR PATHWAY
GO:0051345: positive regulation of hydrolase activity
GO:0043462: regulation of ATPase activity
hsa00270: Cysteine and methionine metabolism
M106: PID HNF3B PATHWAY
0
2
4
6
8
10
12
-log10(P)

**Table S1.** Antibodies were obtained from ebioscience, BD Biosciences, Miltenyi, Beckman-Coulter, Cell Signaling Technology or Biolegend.

| Human Antigen | Clone | Human Antigen/flurorochromes |
|---|---|---|
| 2B4 | 2-69 | 2B4 PE |
| CD107a | eBioH4A3 | CD107a FITC |
| CD14 | MoP9 | CD14 PECF594 |
| CD16 | 3G8 | CD16 BV510 |
| CD160 | BY55 | CD160 AF488 |
| CD19 | HIB19 | CD19 PECF594 |
| CD3 | SK7 | CD3 APC-Cy7 |
| CD3 | SK7 | CD3 BV605 |
| CD39 | TU66 | CD39 BV650 |
| CD4 | RPA.T4 | CD4 PECF594 |
| CD56 | NCAM16.2 | CD56 APC |
| CD56 | AF127H3 | CD56 PE Vio770 |
| CD56 | NCAM 16.2 | CD56 BV786 |
| CD57 | NK-1 | CD57 BV421 |
| CD7 | M-T701 | CD7 BUV395 |
| CD71 | M-A712 | CD71 BV421 |
| CD98 | UM7F8 | CD98 PE |
| DNAM-1 | DX11 | DNAM-1 BV605 |
| EOMES | WD1928 | EOMES PerCP ef710 |
| GRANZYME B | GB11 | GRANZYME B AF700 |
| IFNg | 4SB3 | IFN-g PE |
| KLRG1 | 2F1 | KLRG1 BV510 |
| LAG-3 | 3DS223H | LAG-3 PE Cy7 |
| MIP1b | D21.13.51 | MIP1b v450 |
| NKG2A | REA110 | NKG2A PE Vio770 |
| NKG2C | 134591, R&D Systems | NKG2C PE |
| NKG2D | 1D11 | NKG2D BV650 |
| NKp30 | p30-15 | NKp30 BV510 |
| NKp46 | 9.E2 | NKp46 AF700 |
| pAkt S473 | M89-61 | pAkt PE S473 |
| PD-1 | EH12.1 | PD-1 BV786 |
| PERFORIN | dG9 | PERFORIN BV421 |
| pS6 (pS235/236) | D57.2.2E or D52.2.2E | pS6 (pS235/236) PB |
| pSTAT5 (pY694) | 47/Stat5(pY694) | pSTAT5 PerCPCY5.5 (pY694) |
| T-BET | O4-46 | T-BET BV711 |
| TIGIT | MBSA43 | TIGIT PerCPef710 |
| TIM-3 | 7D3 | TIM-3 BB515 |
| TNFa | mAb 11 | TNFa PE Cy7 |
| TOX | REA473 | TOX PE |
